# Structure of the host cell recognition and penetration machinery of a *Staphylococcus aureus* bacteriophage

**DOI:** 10.1101/721746

**Authors:** James L. Kizziah, Keith A. Manning, Altaira D. Dearborn, Terje Dokland

## Abstract

*Staphylococcus aureus* is a common cause of infections in humans. The emergence of virulent, antibiotic-resistant strains of *S. aureus* is a significant public health concern. Most virulence and resistance factors in *S. aureus* are encoded by mobile genetic elements, and transduction by bacteriophages represents the main mechanism for horizontal gene transfer. The baseplate is a specialized structure at the tip of bacteriophage tails that plays key roles in host recognition, cell wall penetration, and DNA ejection. We have used high-resolution cryo-electron microscopy to determine the structure of the *S. aureus* bacteriophage 80α baseplate at 3.7 Å resolution, allowing atomic models to be built for most of the major tail and baseplate proteins, including two tail fibers, a trimeric receptor binding protein, and part of the tape measure protein. Our structure provides a structural basis for understanding host recognition, cell wall penetration and DNA ejection in viruses infecting Gram-positive bacteria. Comparison to other phages demonstrate the modular design of baseplate proteins, and the adaptations to the host that take place during the evolution of staphylococci and other pathogens.

## INTRODUCTION

*Staphylococcus aureus* is a Gram-positive bacterium and opportunistic pathogen that colonizes the nasal cavities of ≈30% of the population, increasing the risk of pathogenic infection, especially in the clinical setting (*1*). Almost 120,000 cases of *S. aureus* blood stream infections occurred in the United States in 2017, with 20,000 associated deaths (*2*). *S. aureus* resistant to methicillin (MRSA) and other antibiotics have become a major public health concern. Most resistance and virulence factors in *S. aureus* are encoded on mobile genetic elements (MGEs), including bacteriophages (phages) and genomic islands (*3, 4*). Given the rarity of genetic conjugation loci and restriction against transformation, transduction via bacteriophages is considered the main mode of horizontal gene transfer of MGEs in *S. aureus* (*5*). Some staphylococcal bacteriophages carry virulence factor genes in their genomes. Many phages are also capable of transferring unrelated genetic material through generalized transduction. Furthermore, certain “helper” phages are involved in highly specific, high frequency mobilization of *S. aureus* pathogenicity islands (SaPIs) and similar genetic elements (*6, 7*). The underlying mechanisms of horizontal gene transfer and the ability of a particular phage to transfer genetic material to a new host are therefore central to genetic diversification and the emergence of novel pathogenic strains in *S. aureus*.

Bacteriophages of many Firmicutes, including *S. aureus*, interact with wall teichoic acid (WTA), a variable carbohydrate polymer present on the surface of most Gram-positive cells (*8*). Most strains of *S. aureus* have a unique type of WTA that is distinct from that of other (coagulase-negative) staphylococci (CoNS) and generally blocks infection by CoNS-specific phages (*9*). A key determinant of host specificity is provided by the phage tail tip complex (TTC) or more elaborate baseplate, specialized structures at the tips of phage tails that are usually the first point of contact between the phage and its host (*10, 11*). Enzymatic activities associated with TTCs and baseplates degrade the cell wall, ultimately leading to penetration of the plasma membrane and ejection of the encapsidated DNA into the cell. Tailed, double-stranded (ds) DNA phages (order *Caudovirales*) are divided into three families based on the structures of their tails: long and flexuous (*Siphoviridae*); long and contractile (*Myoviridae*); or short (*Podoviridae*) (*12*). TTCs and baseplates from these diverse groups vary greatly in complexity, but tend to share a common set of structural modules, combined with a variable set of host interaction proteins, including tail fibers and receptor binding proteins. The best-described bacteriophage baseplate structures are those of the lactococcal siphoviruses TP901-1, Tuc2009, and p2, and the *Escherichia coli* myovirus T4, for which various structures have been produced (*11, 13*). In contrast, little structural information is available for baseplates from staphylococcal bacteriophages.

Phage 80α is a typical staphylococcal siphovirus with a 43.8 kbp genome and a 63 nm icosahedral capsid, attached to a 190 nm long flexuous tail that is capped by an ornate baseplate (*14–16*). 80α is one of the best-described *S. aureus* phages, and is closely related to ϕETA2 and other phages involved in specifying bacterial pathogenicity (*14*). 80α serves as the mobilizing phage for several SaPIs (*14*). The tail and baseplate gene cluster in 80α includes open reading frames (ORFs) 53–68, which are part of the large late operon (ORFs 40–73) that encodes all structural proteins, the terminase proteins involved in DNA packaging, and the lysis proteins (Fig. 1A, B). Here, we have determined the cryo-EM structure of the 80α baseplate at 3.7 Å resolution, allowing most of the baseplate proteins and part of the tail to be modeled in atomic detail. This is the first high-resolution structure of a baseplate from a staphylococcal siphovirus and of a siphovirus tail. The 80α baseplate structure has several unique features, including the presence of three tail fiber/receptor binding proteins, but also displays striking similarities to other phage baseplates, indicating a mix-and-match strategy for baseplate evolution across the Firmicutes. Our structure contributes to a broader understanding of the diversity of the cell recognition and penetration machinery of siphoviruses infecting Gram-positive bacteria and how these structures access the cell interior and trigger DNA ejection during infection.

**Figure 1.**
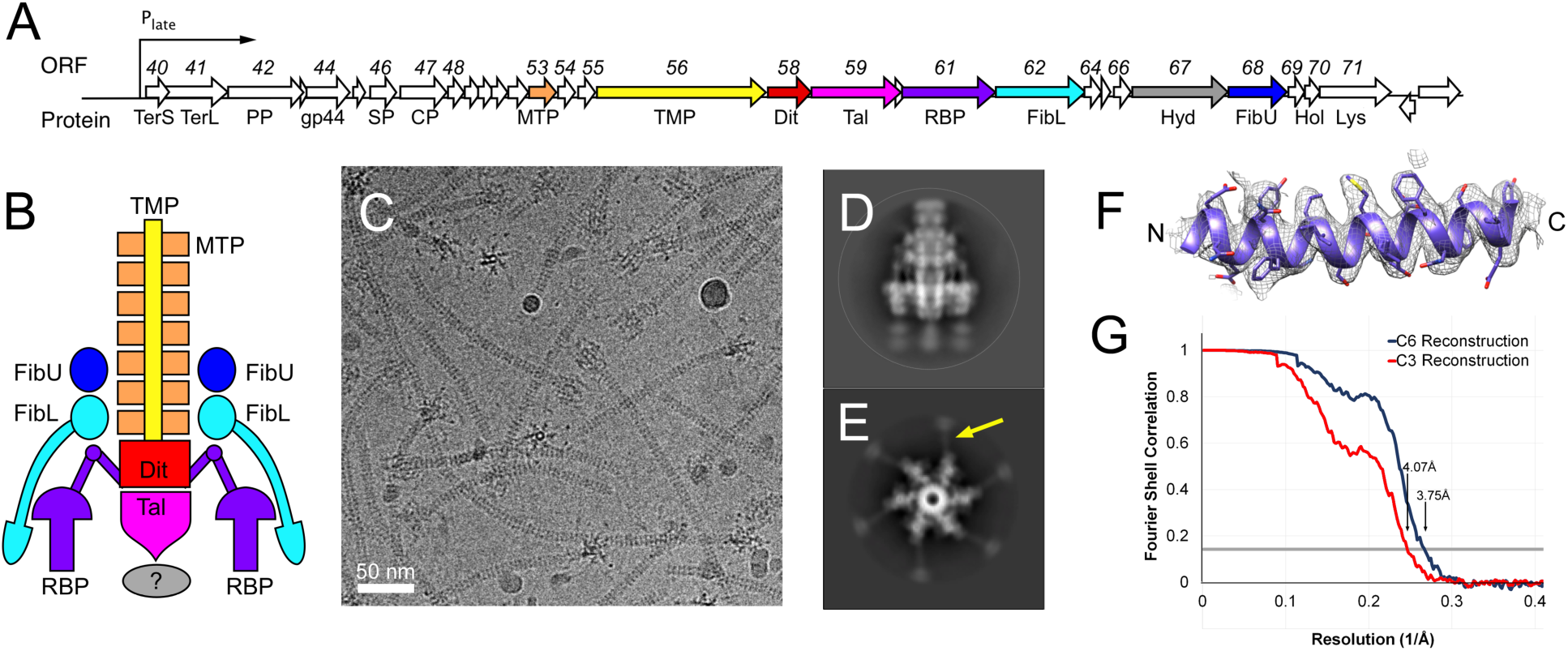
The 80α baseplate. (A) Schematic diagram of the late operon of bacteriophage 80α. ORFs (arrows) corresponding to tail and baseplate proteins are colored tan (MTP), yellow (TMP), red (Dit), pink (Tal), purple (RBP), light blue (FibL), dark blue (FibU), and gray (Hyd). (B) Schematic diagram of the baseplate, colored as in A. The question mark indicates an observed density of uncertain identity, probably corresponding to the Tal CTD. (C) Cryo-electron micrograph of 80α tails collected on a Titan Krios microscope with a DE-20 detector. Scale bar = 50 nm. (D, E) 2D class averages, showing the baseplate in side (D) and top (E) view, viewed perpendicular and parallel to the long axis of the tail, respectively. A thin fiber is indicated with a yellow arrow in E. (F) Representative density from the C6 reconstruction with the corresponding atomic model. (G) Fourier shell correlation curve between two half datasets for the C6 (blue) and C3 (red) reconstructions. The resolution at FSC=0.143 (gray line) is indicated for each.

## RESULTS

### 1. Three-dimensional reconstruction of the 80α baseplate

80α tails were produced using an 80α lysogen (strain ST247) with a deletion of the first 14 amino acids of the scaffolding protein (gp46), which results in failure to form capsids and accumulation of tails in the lysate (Fig. 1C) (*16*). Initial cryo-EM datasets of up to ≈7,000 baseplate particle images were collected with a Philips CM20 microscope on SO-163 film or using an FEI Titan Krios microscope equipped with a Direct Electron DE-20 detector, and reconstructed with C6 symmetry to 11 Å resolution (FSC_0.5_) using EMAN and EMAN2 (*17*). A larger dataset was subsequently collected with the Titan Krios and DE-20 detector, and processed with MotionCor2, Gctf and RELION2 (*18*), yielding a final C6 reconstruction at 3.75 Å resolution (FSC_0.143_) from 74,294 particle images (Fig. 1D– G, Supplementary Figs S1 and S2). A reconstruction of the same data with application of only C3 symmetry was generated with RELION3 (*19*) and reached 4.07 Å resolution (Fig. 1G, Supplementary Figs S1 and S2).

The baseplate is 298 Å wide and 271 Å tall and consists of six peripheral structures surrounding a core that represents a continuation of the tail (Fig. 2A, B). The tail itself is 86 Å wide and consists of ≈39 hexameric rings with an inter-ring spacing of 41 Å, enclosing a 42 Å wide lumen (Fig. 2B). Three of the tail rings are embedded within the baseplate itself. The peripheral structures consist of two large rings surrounding the embedded part of the tail, and six trefoiled club-shaped features, about 85 Å wide and 124 Å long, connected to the baseplate core via six kinked stems. At lower cutoff level, six fibers can be seen to extend from the lower ring and wrap around the outside of the baseplate, terminating in globular densities near the baseplate tip (Fig. 2C). At this cutoff level, at least seven tail rings can be observed, and a rod-like protrusion extends from the tip of the baseplate. Additional, noisy densities, disconnected from the rest of the baseplate, appear below this protruding rod. 2D class averages that represent end-on views of the baseplate (Fig. 1E) suggested the presence of six thin fibers spread out radially in the plane of the ice, but these features were not observed in the 3D reconstruction.

**Figure 2.**
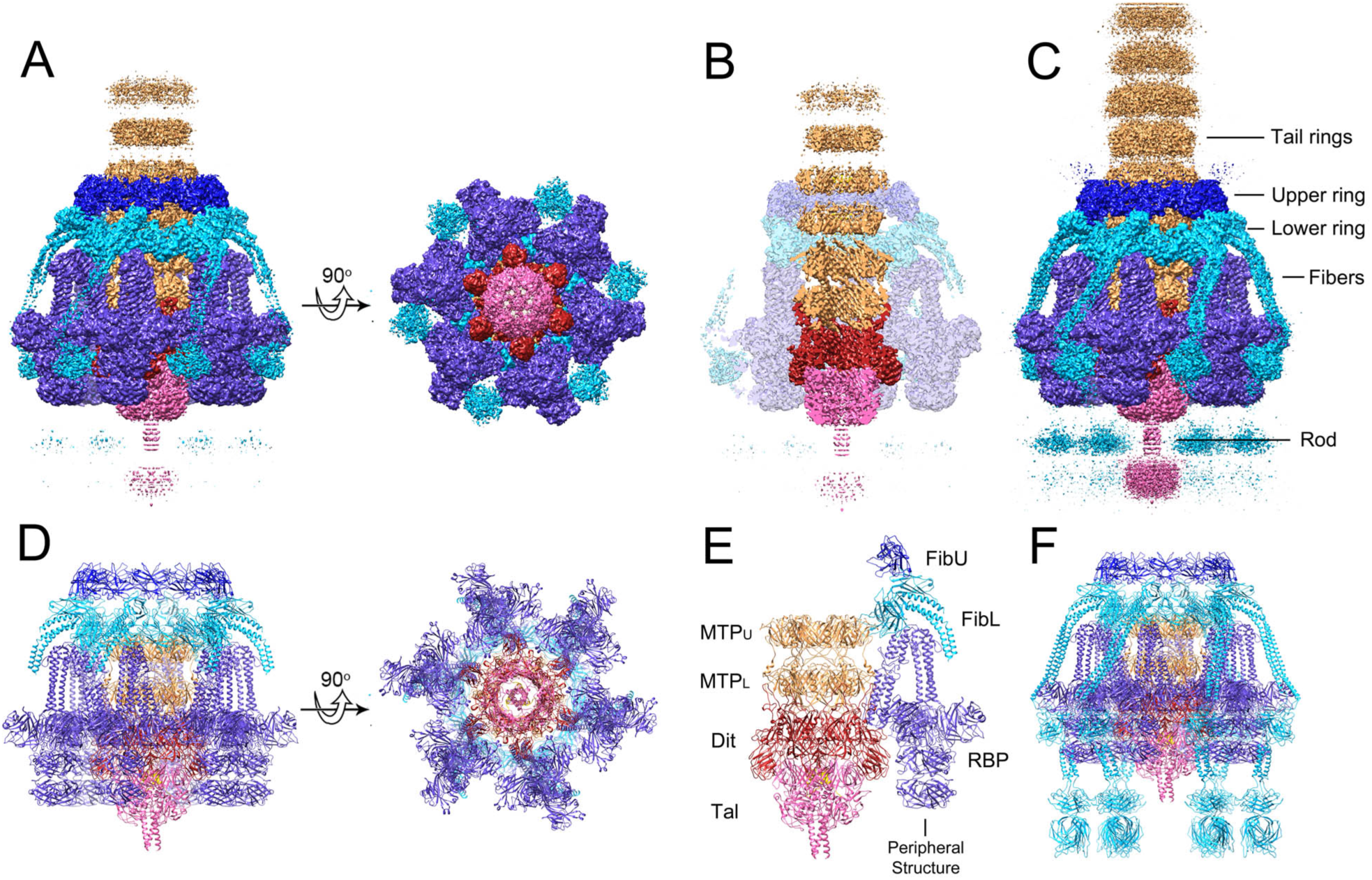
Reconstruction and modeling of the 80α baseplate. (A) Isosurface representation of the 80α baseplate C6 reconstruction viewed from the side and bottom, segmented and color-coded according to the scheme in Fig 1. (B) Cutaway side view, showing the inside of the tail. Peripheral structures are shown as a transparent surface. (C) Side view at lower density cutoff level, showing the tail fibers, tail rings and rod density more clearly. (D) Ribbon representation of the atomic model of the entire baseplate, viewed form the side and bottom and colored as above. (E) Model with all except one peripheral structure removed to show MTP_U_, MTP_L_, Dit and Tal underneath. (F) Complete baseplate model showing the full-length FibL pseudo-atomic model.

### 2. Model building of baseplate proteins

In spite of limited sequence homology, some of the tail and baseplate genes could be assigned based on similarity to other phages, including those encoding the major tail protein (MTP), gene product (gp) 53 (ORF53), and the tape measure protein (TMP, gp57), which is generally the largest ORF in siphovirus genomes (Fig. 1A, Table 1). Other ORFs could be assigned by comparison with the closely related phage ϕ11 (*20*), for which HHpred analysis (*21*) had identified homologs in the PDB structure database. These ORFs include the distal tail protein (Dit, gp58), which comprises the baseplate “hub”, the tail-associated lysin (Tal, gp59), comprising the tip of the tail, and the receptor binding protein (RBP, gp61) (Fig. 1A, B; Table 1). The N-terminal part of ϕ11 gp54, corresponding to 80α gp62, was shown to have some similarity with upper baseplate protein (BppU) of lactococcal phage TP901-1 (*20, 22*). The crystal structure of ϕ11 RBP (gp45), which is 97% identical to the 80α RBP, was recently determined (*23*). Crystal structures have also been determined for SPP1 Dit (*24*) and the baseplates of lactococcal phages p2 and TP901-1, which include Dit, Tal, BppU, and receptor binding proteins that are not similar to the ϕ11 RBP (*22, 25*).

**Table 1.** List of 80α tail and baseplate proteins, showing ORF numbers in 80α and corresponding numbers in ϕ11, protein name, number of amino acid residues and calculated molecular weight, number of copies per particle, and a description of protein functions and location in the virion.

| 80α<br>ORF | ϕ11<br>ORF | Protein<br>name | Residues | MW<br>(kDa) | Copies | Function and location |
| --- | --- | --- | --- | --- | --- | --- |
| 53 | 39 | MTP | 193 | 21.5 | ≈234 | Major tail protein; forms ≈39 hexameric rings |
| 57 | 42 | TMP | 1154 | 125.8 | 3 | Tape measure protein; center of tail; C-terminus inside Tal |
| 58 | 43 | Dit | 315 | 37.1 | 6 | Baseplate hub, hexameric ring |
| 59 | 44 | Tal | 633 | 71.0 | 3 | Baseplate tip; trimeric ring; C-terminal hydrolase domain |
| 61 | 45 | RBP | 636 | 73.7 | 18 | Receptor (WTA) binding protein; 6 trimers |
| 62 | 54 | FibL | 607 | 66.8 | 18 | Lower fiber, α-helical coiled-coil; 6 trimers |
| 67 | 49 | Hyd | 632 | 72.0 | unknown | Peptidoglycan hydrolase; not seen in map |
| 68 | 50 | FibU | 390 | 43.8 | 12? | Upper fiber, collagen-like; 6 dimers seen in map |

We carried out HHpred analysis on 80α proteins gp53, gp57, gp58, gp59, gp61, gp62, gp67 and gp68, which identified additional homologies compared to ϕ11 due to server improvements (*21*) and the presence of more structures in the database (Table 2). Most importantly, gp62 was identified as a tail fiber (FibL) with coiled-coil structure, while gp68 was predicted as a collagen-like fiber protein (FibU). gp67 encodes a predicted cell wall hydrolase (Hyd) that is 97% identical to ϕ11 gp49, which was previously shown to exhibit muralytic activity (*26*). Gp67 was the only putative baseplate protein that could not be identified in the baseplate reconstruction, presumably because it is disordered and/or present at low occupancy, consistent with previous MS analysis (*27*).

**Table 2.**
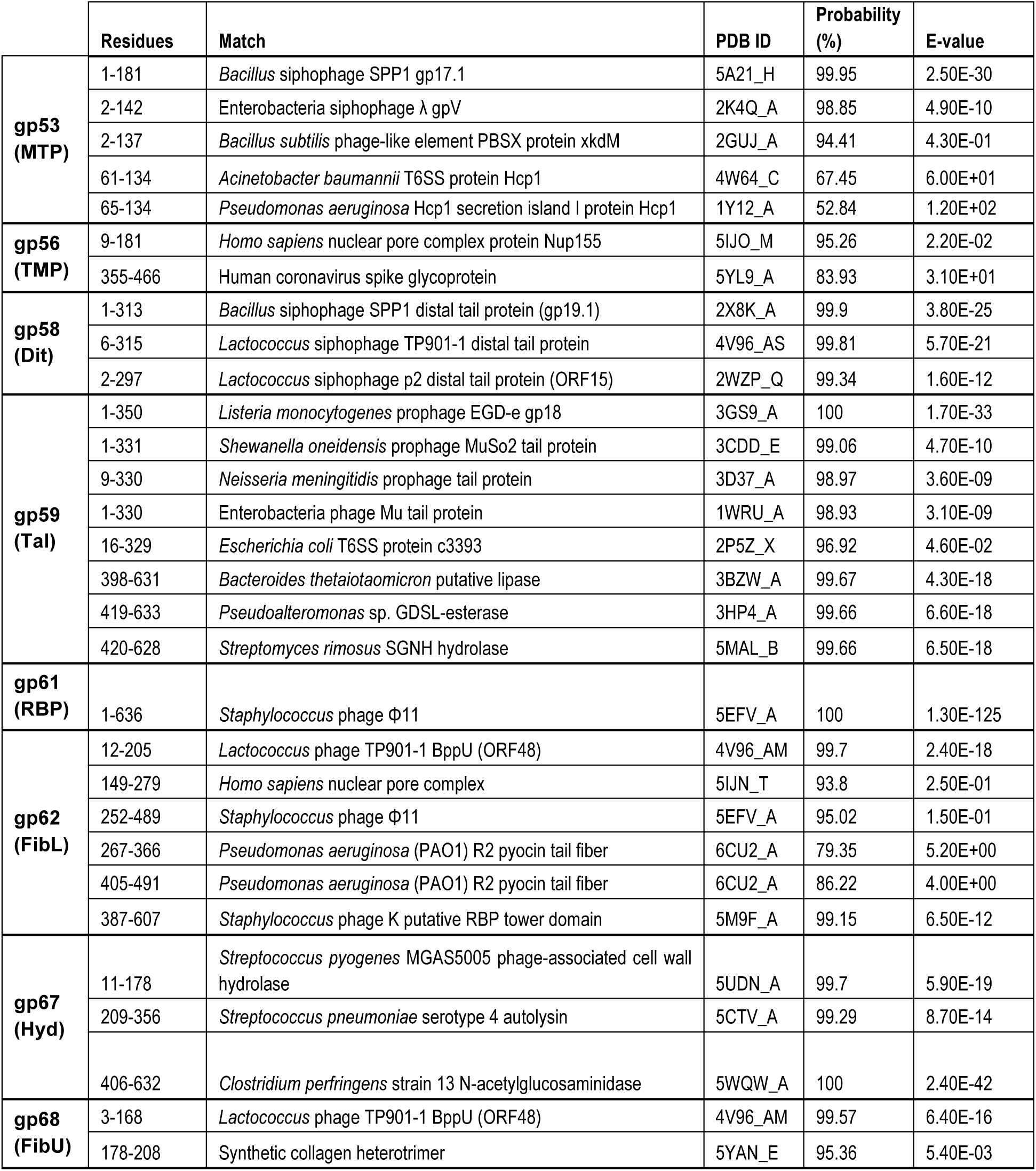
HHpred analysis of 80α baseplate protein sequences. The most relevant hits are shown for each protein, with matched residues, PDB ID and chain identifier, HHpred probability (%) and E-value.

Initial atomic models for MTP (gp53), Dit (gp58), and parts of Tal (gp59), FibL (gp62), FibU (gp68) and TMP (gp57) were generated in I-TASSER (*28*). A model for RBP (gp61) was generated by manually replacing the 16 residues that differ between gp61 and the ϕ11 gp45 crystal structure (*23*). These starting models were fitted into either the C6 or C3 baseplate reconstruction, guided by the matching of large amino acid side chains, followed by cycles of manual model building and real-space refinement in Phenix (*29*). The RBP, FibL and FibU models were further improved by refinement into the focused reconstructions (see Methods). In total, 72 polypeptides comprising 7 different proteins (Table 1) were included in the final model (Fig. 2D–F). The FSC curves between the final models and the corresponding reconstructions are shown in Supplementary Fig. S1.

### 3. The major tail protein (MTP, gp53)

The major tail proteins of siphoviruses, tail tube proteins of myoviruses, and the Hcp1 tube-forming proteins of type 6 secretion systems (T6SS) share a common fold, known as the tail tube fold (*30*). In addition, the tail tube fold has been identified in various baseplate proteins, tail terminator proteins, and the T4 capsid assembly protease, reflecting their evolutionary relationship (*31*). HHpred analysis of 80α gp53 yielded >98% probability matches to major tail proteins of several siphoviruses, including SPP1 gp17.1 and λ gpV (*32, 33*), and lower probability matches to two T6SS tube-forming proteins (Table 2).

Six copies of MTP were built into each of the two most well defined tail rings closest to the baseplate. Of the 193 residues of MTP, residues 4-169 were modeled in the lower ring (denoted MTP_L_) and 4-150 into the upper ring (MTP_U_) (Fig. 2E, 3A). Each MTP subunit includes two four-stranded β-sheets folded into a β-sandwich and flanked by an α-helix (Fig. 3A). In a complete tail ring, one of these β-sheets forms a 24-stranded, highly negatively charged β-barrel—42 Å in diameter— lining the tail lumen, presumably providing a slippery tube for the DNA during infection (Fig. 2B, 3B). This luminal β-sheet contains an extended insertion loop between β2 and β3 (the “stacking loop”) that reaches across the ≈3 Å space that separates tail rings and inserts into a pocket between two subunits in the adjacent ring (Fig. 3A). The equivalent loop in the major tail proteins of phages λ, T5 and SPP1 was shown to play an essential role in tail polymerization (*32–34*).

**Figure 3.**
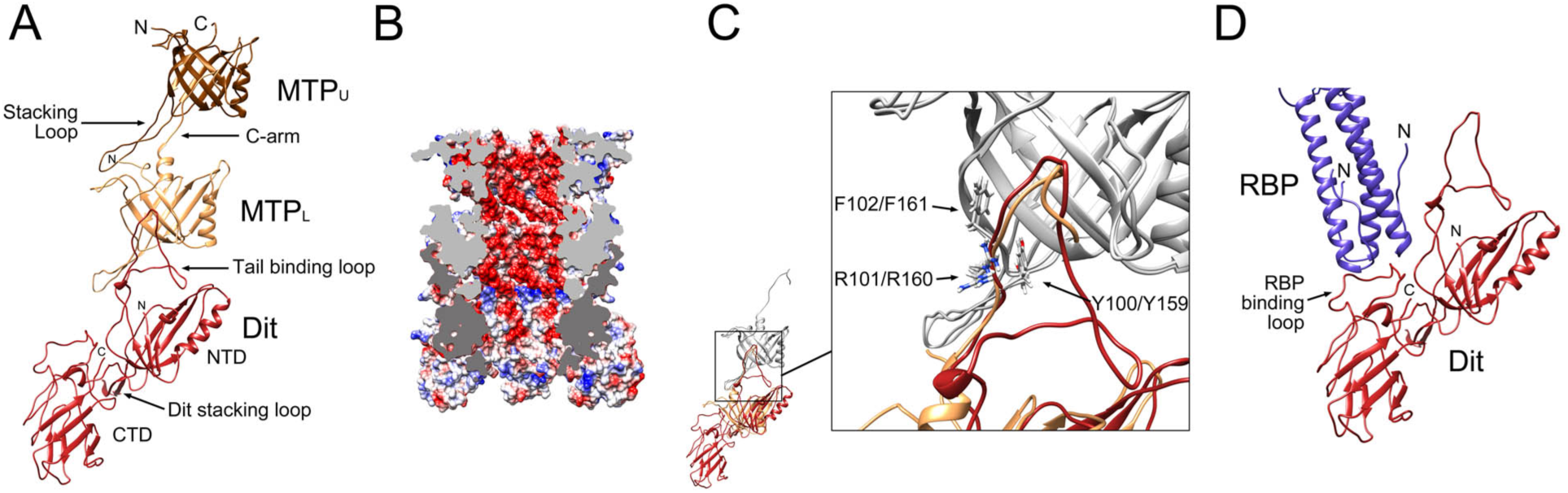
The baseplate core. (A) One asymmetric unit of the sixfold symmetric baseplate core, showing MTP_U_ (brown), MTP_L_ (tan) and Dit (red). The stacking loops of MTP and Dit, the C-arm of MTP_L_ and the Dit tail binding loop are indicated. (B) Cutaway electrostatic surface showing the charge distribution inside the tail, colored from red (negative) to blue (positive). The cut surfaces of MTP and Dit are colored light and dark gray, respectively. (C) Superposition of MTP_U_ onto MTP_L_ (gray). MTP_L_ (tan) shifted by the same amount then superimposes on Dit (red). The expanded view shows the superposition of the Dit tail binding loop (red) with the MTP C-arm (tan). The side chains of the triplet of residues conserved between the C-arm of MTP (Y159,R160,F161) and the tail binding loop in Dit (Y100,R101,F102) are shown in stick representation and labeled. (D) Detail of the interaction between Dit (red) and the RBP coiled-coil stem region (purple), showing the RBP binding loop in the Dit CTD.

An additional inter-ring contact is mediated by the C-terminus of MTP (the “C-arm”), which extends in the opposite direction to the stacking loop and fills a crevice on the exterior surface of the adjacent subunit (Fig. 3A). No equivalent contact has been seen in other systems. The major tail proteins of phages λ, SPP1 and T5 instead have an additional C-terminal immunoglobulin (Ig)-like domain (*32– 34*).

### 4. Distal tail protein (Dit, gp58)

The “distal tail protein” (Dit) constitutes the hub of the baseplate, and is highly conserved between TTCs and baseplates of siphoviruses (*31*). HHpred matched 80α gp58 to the Dit proteins of several phages with ≥99.5% probability, including SPP1 gp19.1, TP901-1 ORF46 and p2 ORF15 (Table 2) (*22, 24, 25*). Dit forms a connection between the sixfold symmetric tail and the threefold symmetric tail tip (Tal, see below), and provides an attachment site for the six RBP trimers (Fig. 2D–F).

Six copies of Dit were built into the hexameric ring of density in the baseplate core just below MTP_L_ (Fig. 2E, 3A). Dit consists of two domains: an N-terminal domain (NTD, residues 1–166) with a tail tube fold, and a C-terminal domain (CTD, residues 167–315) with a galectin-like fold (Fig. 3A). The NTD is similar to MTP and includes a stacking loop—previously described as the “belt extension” in SPP1 (*24*)—that interacts with Tal (Fig. 3A). The interior surface of the Dit NTD forms a negatively charged β-barrel that constitutes a continuation of the tail lumen (Fig. 2B, 3B). The Dit NTD lacks a C-arm equivalent to that of MTP; instead, an extended insertion loop (the “tail binding loop”) between β4 and β5 of Dit fits into the same crevice on MTP_L_ that the C-arm of MTP_L_ occupies on MTP_U_ (Fig. 3A, C). This topologically distinct Dit loop contains a triplet of residues (Y100, R101, F102) that matches a corresponding triplet in the MTP C-arm (Y159, R160, F161), a striking case of convergent evolution (Fig. 3C).

The Dit CTD is also conserved between 80α and SPP1, TP901-1 and p2 (*22, 24, 25*). However, the 80α Dit has an insertion (the “RBP binding loop”, residues 182-197) that is found in p2, but not in SPP1 and TP901-1. As in p2, this loop serves as the attachment site for the RBP (Fig. 3D). In TP901-1 Dit, which lacks this loop, peripheral structures (BppU) are instead bound to the Dit NTD, whereas SPP1 lacks peripheral structures.

### 5. Tail-associated lysin (Tal, gp59) and tape measure protein (TMP, gp57)

HHpred analysis matched the NTD of 80α gp59, consisting approximately of residues 1–350, to baseplate-related proteins from several phages that share similarity with the gp27 hub protein that constitutes part of the cell-piercing apparatus of *E. coli* phage T4 (Table 2) (*31*). These trimeric proteins contain two tail tube folds, resulting in a quasi-C6 structure that binds Dit to cap off the baseplate. The CTD of gp59, comprising residues 390–633, matched various esterases and lipases of the GDSL-hydrolase family and the SGNH-hydrolase subfamily (*20*) (Table 2), suggesting a role in cell wall degradation. For consistency with what was previously proposed for ϕ11 (*20*), we will refer to this protein as Tal, for “tail-associated lysin,” although the enzymatic activity of this protein has not yet been established.

The Tal NTD was initially fitted as a C3 trimer into the cap-like structure at the bottom of the baseplate, directly below the Dit hexamer in the C6 symmetric reconstruction (Fig. 2A). Due to the quasi-sixfold nature of the Tal trimer, it matched quite well overall, but much of the density was blurred due to the superposition of non-identical Tal orientations. At the C-terminal end of the Tal NTD, the density formed a thin rod that connected to a noisy globular density, neither of which could be interpreted in the C6 reconstruction. To obtain a reconstruction with only threefold symmetry imposed, we first used a masked reconstruction to subtract all density not associated with Tal from the baseplate images. These signal-subtracted images were then subjected to symmetry expansion and 3D classification to group together consistently oriented Tal trimers within the larger C3 asymmetric unit. These orientations were used to generate a reconstruction of the whole baseplate with C3 symmetry, which reached a resolution of 4.07 Å (Fig. 1G). In this reconstruction, the density corresponding to the cap and the rod was well resolved (Fig. 4A–C) and allowed modeling of residues 2–388 of Tal (Fig. 4D, E). The disorganized density beyond the rod was presumed to corresponds to the Tal CTD (residues 390 to 607), but was not interpretable even in the C3 reconstruction (Fig. 4C).

**Figure 4.**
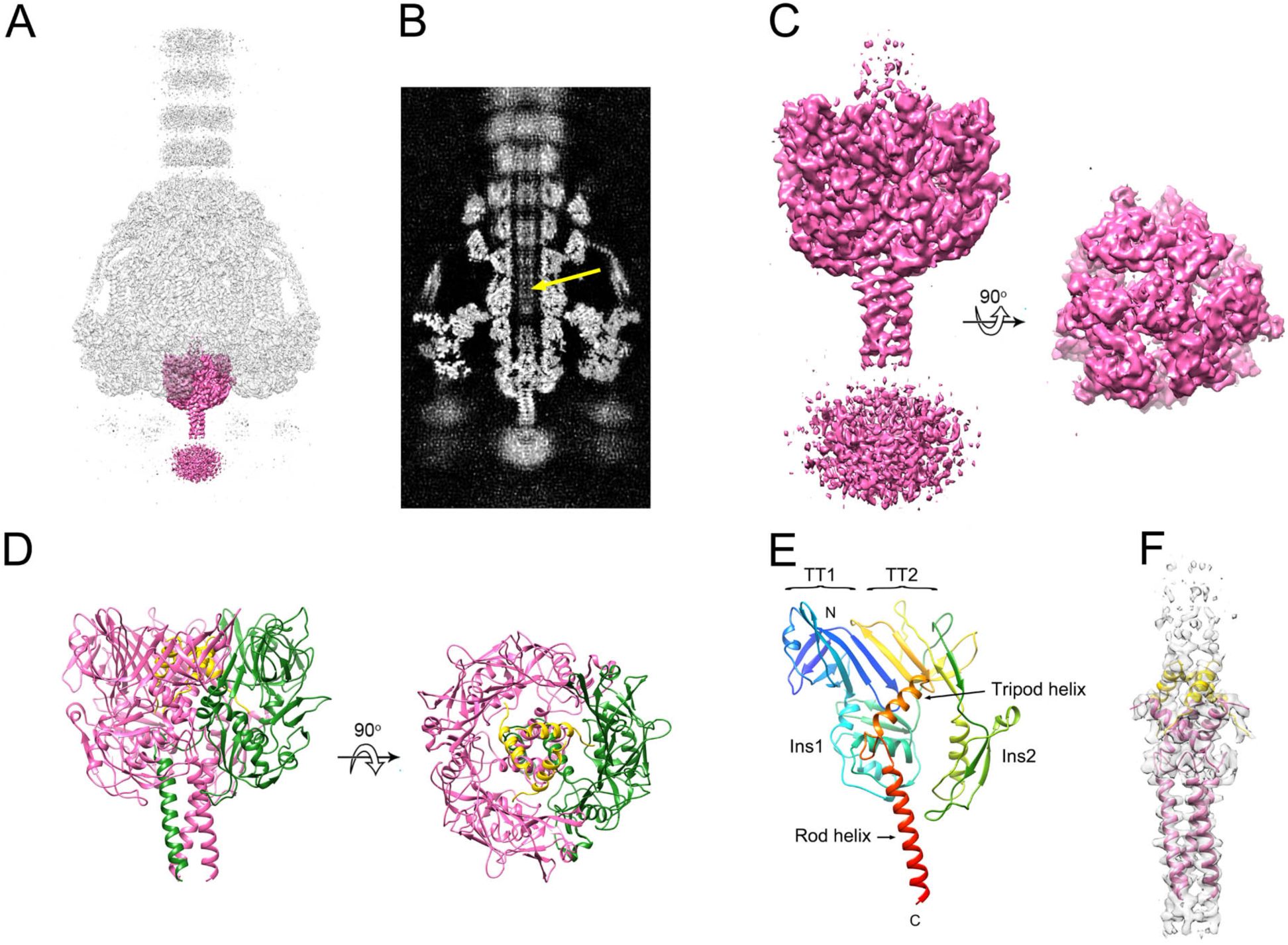
The baseplate tip. (A) Isosurface representation of the C3 symmetric baseplate reconstruction. The density corresponding to the Tal trimer is shown in pink, with the rest of the reconstruction in transparent gray. (B) Central section through C3 reconstruction (gray scale). Density inside the lumen presumably corresponding to TMP is indicated with the yellow arrow. (C) The Tal density alone, viewed from the side and bottom. The disordered density at the bottom in the side view is assumed to be the Tal CTD and was removed from the bottom view for clarity. (D) Ribbon representation of the atomic model of Tal viewed from the side and bottom. One subunit is colored green. The TMP model is shown in yellow. (E) Tal subunit, colored in rainbow colors from blue (N-terminus) to red (C-terminus). Relevant structural elements are labeled. (F) Density corresponding to the central part of Tal with the atomic models of Tal and TMP shown in pink and yellow, respectively.

The Tal NTD (residues 1-331) consists of two tail tube domains (TT1 and TT2) that form a quasi-sixfold adapter to the Dit hexamer (Fig. 4D). The two domains merge into a continuous 8-stranded β sheet on one side (Fig. 4E). The two domains are separated by an insertion (Ins1) with a fold related to the type 3 secretion system protein EscC (PDB ID: 3GR5). The second tail tube domain has an additional domain (Ins2) inserted in a position that is topologically equivalent to the β2-β3 stacking loop in MTP (Fig. 4E). Residues 335-348 form an α-helix that extends into the tail lumen. In the Tal trimer, these helices form a twisted tripod that occludes the opening of the lumen (Fig. 4D, F). Residues 362-389 from the three subunits make up an α-helical coiled-coil that forms the rod and extends to the unresolved CTD (Fig. 4D, F).

After accounting for the plug and rod helices, there were three additional, well-ordered α-helical densities forming a second tripod interlaced with that of the Tal plug (Fig. 4D, F). Although not conclusively determined, this density matched the C-terminal residues 1135-1154 of TMP. TMP is the only protein known to fill the tail channel and the localization of its C-terminus at the baseplate agrees with previous research on other phages (*35*). Additional density attributable to TMP could be seen inside the tail lumen, but was not interpretable in terms of the atomic structure (Fig. 4B).

### 6. Receptor binding protein (RBP, gp61)

80α gp61 is 97% identical in amino acid sequence to gp45 of the closely related phage ϕ11, for which the crystal structure was previously determined (*23*). Gp45 was previously identified as the primary receptor binding protein for ϕ11, with binding affinity for WTA, and was thus denoted RBP (*20*). As expected, the 80α RBP is very similar to ϕ11 gp45 (*23*). RBP is a trimeric protein consisting of four domains: an N-terminal α-helical coiled-coil “stem” domain (residues 1–142) that is folded by 155° at a central hinge; a five-bladed β-propeller “platform” domain (residues 143–442) that was previously suggested to bind to the GlcNAc residues of WTA on the host cell surface (*20, 23*); and two highly similar C-terminal “tower” domains (residues 443–636), each consisting of a bent β-sheet superposed by an α-helix (Fig. 5A). Six such trimers surround the baseplate core, making up the bulk of the observed peripheral structures (Fig. 2D). The greatest difference between the 80α and ϕ11 RBPs is in the orientation of the stem domains relative to the platform and tower domains (Fig. 5B). As noted above, the N-termini of the coiled-coil domains from each RBP trimer connect to the RBP binding loop of Dit (Fig. 3D). The RBP hinge interacts with a ring formed by the FibL trimers (Fig. 2D, see below). Deletion of ORF61 led to a complete loss of peripheral structures and of infectivity, although tails still assembled normally. (Supplementary Fig. S3)

**Figure 5.**
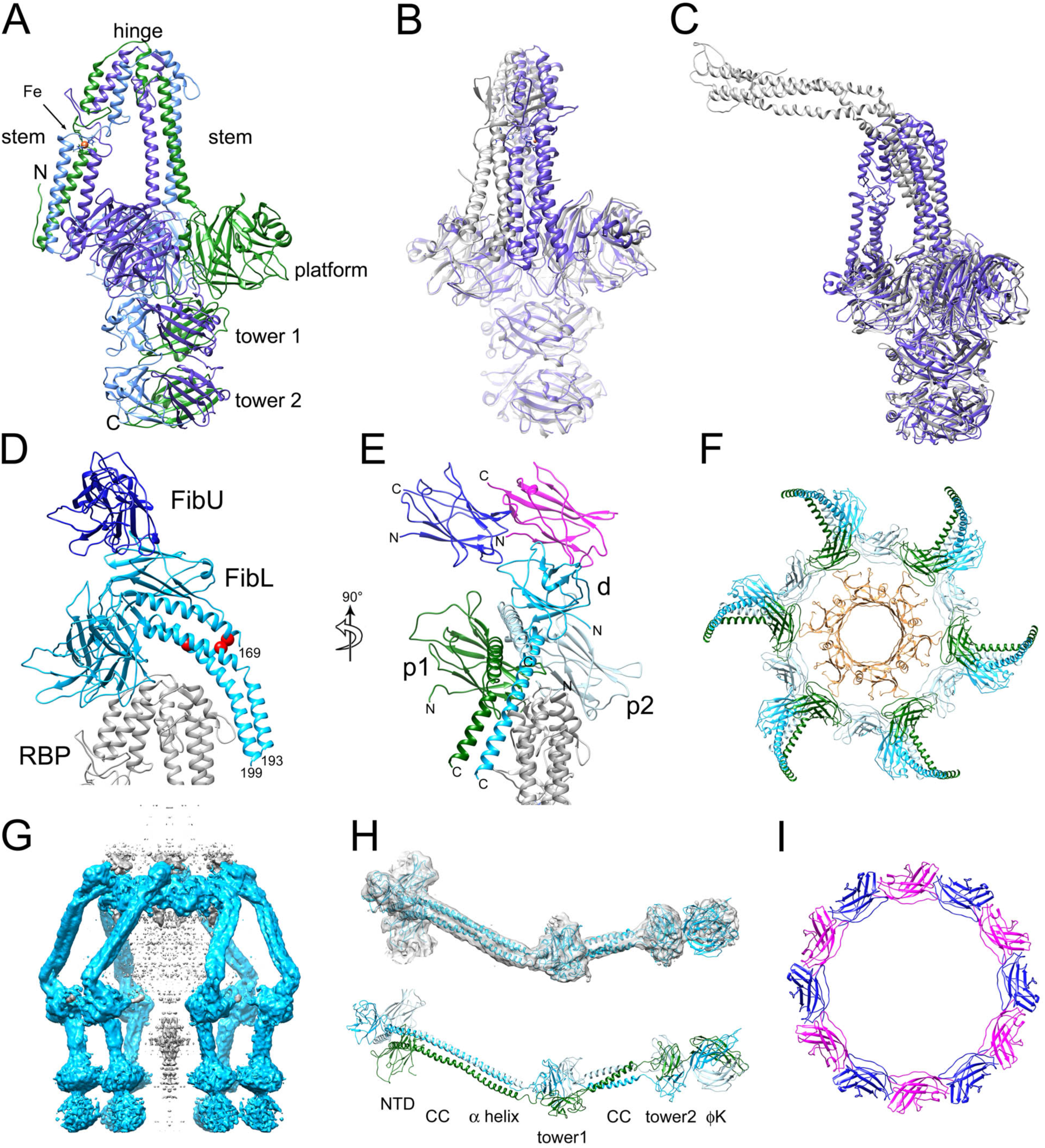
The peripheral structures. (A) Atomic model of the 80α RBP homotrimer. The three subunits are colored purple, blue and green. The stem, hinge, platform and two tower domains are indicated. An Fe atom (orange) coordinated by six His residues in the stem domain is shown. (B) Superposition of 80α (purple) and ϕ11 (gray) RBPs, aligned by the tower and platform domains and rotated to emphasize the differing orientations of the stem domains. (C) Superposition of 80α RBP (purple) with the tail fiber (gp17) of P68 (gray), aligned by the tower domains. (D) Ribbon representation showing FibL (light blue), FibU (dark blue) and part of the RBP stem (gray). The position of residue K166 in the three FibL subunits is shown as red balls to indicate the relative shift of the three α-helices in the coiled coil. The C-terminal residue in each FibL subunit is numbered. (E) Rotated view of D with the three FibL subunits colored green (p1), pale blue (p2) and blue (d). The two FibU subunits are in dark blue and magenta. RBP is gray. (F) Top view of baseplate showing the FibL ring octadecamer (colored as in E) and the MTP hexamer underneath (tan). (G) Isosurface of a reconstruction made from signal-subtracted images excluding density corresponding to MTP, Dit, Tal and RBP. FibL density is blue. Additional density in the center (gray) probably corresponds to TMP. (H) Model for the complete FibL protein trimer built into the density from G. The lower panel is colored as in E, with structural elements labeled (CC, coiled coil; ϕK is the phage K gp68-like domain). (I) Top view of the dodecameric FibU ring, colored as in E.

The structure of RBP is remarkably similar to the “tail fiber” (gp17) of the distantly related staphylococcal podovirus P68 (*36*), presumably reflecting specificity for *S. aureus*. However, in P68 gp17, the stem domain is less bent at the hinge compared to 80α (66° vs. 155°) (Fig. 5C). In addition, since P68 lacks a Dit equivalent, the gp17 stem domain is instead attached to the dodecameric portal protein. Consequently, P68 has twelve gp17 trimers, compared to the six RBP trimers in 80α.

### 7. Lower tail fiber (FibL, gp62)

Gp62 is highly modular, with HHpred matches to several distinct phage protein structures (Table 2). The first 205 residues matched the N-terminal Ig-like domain and α-helical coiled-coil domain of TP901-1 BppU (*22*). The central part of gp62 (residues 146-279) matched several α-helical coiled-coil structures at lower probability (≤93.8%; Table 2). Residues 252–489 were matched to the tower domains from ϕ11 RBP, with lower probability matches of residues 267–366 and 405–491 to a similar domain from a *Pseudomonas aeruginosa* R2 pyocin fiber protein (Table 2). The C-terminal residues of FibL (387-607) matched the two C-terminal domains of the gp68 receptor binding protein from *S. aureus* phage K (*37*), which comprise an RBP-like tower domain and a distinct β-sandwich domain (PDB ID: 5M9F). Because of the fibrous coiled-coil domain and its formation of the lower of the two exterior baseplate rings, the protein was designated FibL, for lower tail fiber.

Six trimers of the N-terminal Ig-like domain of FibL were fitted into the lower of the external rings surrounding the baseplate (Fig. 2D–F, 5D–F). The two proximal subunits (p1 and p2) from each trimer contribute to the ring itself, while the third, distal, subunit (d) is attached on the outside of the ring. Each subunit in the trimer contributes an α-helix to the subsequent coiled-coil domain that extends to residue 169 for subunits p2 that contributes the top α-helix, and to 193 or 199 for the p1 and d subunits, respectively (Fig. 5E). This organization is similar to that of the BppU protein of TP901-1, except that in the lactococcal phages, one of the α-helices in the coiled coil originates from a subunit in the adjacent trimer (*22*). Due to the differing disposition of the N-terminal domains, the residues of the α-helical domains are shifted relative to one another, inducing asymmetry within the coiled coil (Fig. 5D).

The FibL ring is not closely associated with either the Dit or MTP hexamers. Although residues 11 and 12 of one of the FibL subunits approaches the β4-β5 loop of MTP_U_ by ≈4 Å, most of MTP_U_ is separated from the FibL ring by ≈15-20 Å (Fig. 5F). However, the bottom of the Ig-like fold of FibL associates closely with the hinge domain of the RBP stem region (Fig. 5D,E). This is consistent with the observation that deletion of the gene encoding FibL (ORF62) led to a complete loss of peripheral structures, including RBP (Supplementary Figure. S3), suggesting that the interaction between FibL and RBP is co-stabilizing. A similar phenotype was observed when the corresponding gene (ORF54) was deleted in ϕ11 (*38*).

Beyond residue ≈180, the density for the FibL coiled-coil and C-terminal domains was attenuated, likely due to flexibility in the coiled-coil domain (Fig. 2A). At lower cutoff level, however, FibL density could be seen to wrap around the peripheral structures (Fig. 2C). A new reconstruction was made from modified images created by subtracting the density corresponding to the fitted portions of the baseplate (Fig. 5G). This reconstruction reached 5.0 Å resolution (Supplementary Figs S1 and S2). In this map, FibL could be seen as an elongated structure with a globular “knee”, extending beyond the tips of the RBP trimers, terminating in two additional globular densities. While this map was not fully interpretable in terms of the atomic structure of the proteins, models for the remaining domains of FibL based on the HHpred predictions could be rigid body fitted into the density and connected to provide a complete pseudo-atomic model for FibL (Fig. 2F, 5H). In this model, the three coiled-coil α-helices continue with no twist between residues 180–238. This is followed by an RBP-like tower domain that was fitted into the knee in the middle of the fiber, followed by another, shorter coiled-coil region. At the C-terminus of FibL, the predicted phage K-like tower and β-sandwich domains matched the disordered densities at the bottom of the baseplate (Fig. 5G, H).

### 8. Upper tail fiber (FibU, gp68)

Like FibL, gp68 was also predicted to include a BppU-like N-terminal Ig-like domain (99.6% probability; Table 2). However, gp68 lacks the α-helical coiled coil portion of FibL. Instead, residues 178-208 were predicted to match various collagen-like triple helix structures (≤95.4% probability), suggesting that the protein forms a collagen-like fiber. The remaining 180 residues did not match any known structure.

An I-TASSER model for the gp68 NTD matched the density corresponding to the upper external ring that makes up the peripheral structures of the baseplate. We designated this protein FibU, for upper fiber protein. Despite the similarity between the NTDs of FibU and FibL (21% sequence identity), the two proteins could be unambiguously distinguished. Based on its predicted collagen-like domain and similarity to FibL, FibU was expected to form trimers. However, there was only density for twelve copies of FibU in the upper ring, equivalent to the proximal subunits of FibL (Fig. 5E, I). Thus, unless there is a third, distal subunit that is so flexible that it cannot be observed in the density, FibU may actually form dimers in spite of its predicted collagen-like domain. Like FibL, FibU does not make close contacts with either MTP or Dit, but both subunits contact the distal subunit of FibL (Fig. 5D, E). FibU was not essential for infectivity, and deletion of the ORF68 gene did not affect the structure of the rest of the baseplate (Supplementary Fig. S3).

No density was observed for the predicted fibrous portion of FibU beyond residue 123 (Fig. 2C). However, 2D class averages representing top views of the baseplate showed six thin fibers extending 9 nm radially from the baseplate and terminating in a 4 nm knob (Fig. 1E). These features might correspond to FibU or FibL fibers interacting with the air-water interface when the baseplate is oriented in the plane of the ice. When the baseplate is oriented laterally (Fig. 1D), the FibU fibers might be disordered and thus not observed in the reconstruction.

## DISCUSSION

We have determined the structure of the bacteriophage 80α baseplate at 3.7 Å resolution, allowing atomic models to be built for the major tail protein, part of the tape measure protein, and all of the major baseplate-associated proteins except gp67, the predicted cell wall hydrolase, the C-terminal domain of Tal, and the C-terminal part of FibU (gp68). This is the first high-resolution structure of a baseplate from a staphylococcal siphovirus. While the 80α baseplate shares many features with other bacteriophages, the structure is unlike those of the well-described baseplates of *E. coli* phage T4 and lactococcal phages p2 and TP901-1 (*11, 13*). In particular, the presence of three separate tail fiber/receptor binding proteins and their organization into external rings surrounding the tail are unique features of the 80α baseplate.

The Dit hexamer constitutes the hub of the baseplate, forming an adaptor between the sixfold symmetric tail and the threefold symmetric Tal protein, as well as the attachment point for the six RBP trimers. Our identification of the C-terminus of TMP associated with Tal supports the assumption that tail assembly starts from a nucleus containing Dit, Tal, and TMP. The first MTP ring is added through interactions involving the MTP stacking loop and the Dit tail binding loop (Fig. 3). Additional MTP subunits are added through interactions of stacking loops and C-arms until the full length of the TMP is reached (*10*). The peripheral structures are not required for this process, since deletions of the genes encoding RBP, FibL or FibU had no effect on tail formation (Supplementary Fig. S3). The addition of RBP and FibL to the hub appears to be interdependent, whereas FibU is probably the last protein to be added to the baseplate.

RBP (gp62) is the main receptor binding protein of 80α, and is essential for infectivity. The virtually identical RBP of ϕ11 was shown to bind wall teichoic acid (WTA) (*20*), a polymer present in the cell wall of most Gram-positive organisms. WTA serves as receptor for many phages, but is highly variable between different species. Most strains of *S. aureus* have WTA that is distinct from that of coagulase-negative staphylococci (CoNS) and is made from ribitol phosphate repeating units modified by GlcNAc (*8, 9*). However, some pathogenic *S. aureus* strains express an altered form of WTA, which results in altered phage susceptibility and immune system evasion (*9*).

Host specificity is reflected in the structure of the receptor binding structures of the infecting phages. The distantly related *S. aureus*-infecting podovirus P68 has a “tail fiber” protein (gp17) that is remarkably similar to the 80α RBP (*36*) (25% sequence identity), suggesting that this structure is conserved among phages that infect *S. aureus*. In contrast, phage Andhra, which is structurally very similar to P68, but infects *Staphylococcus epidermidis* (*39*), has a completely different predicted RBP structure based on a β-helical architecture. *S. epidermidis*, a CoNS species, has a different type of WTA from *S. aureus*, which could explain this difference. Similarly, *S. aureus* phage ϕ187 infects a lineage of *S. aureus* (ST395) that has a WTA more similar to that of CoNS, and is refractory to infection by typical *S. aureus* phages like 80α (*9*). ϕ187 lacks an RBP-like protein, but has a tail fiber protein (gp24) that, by HHpred analysis, resembles parts of FibL, including an N-terminal Ig-like domain, a coiled-coil domain, a tower domain, and a C-terminal 7-stranded β-sandwich domain related to the receptor binding domain of the phage K tail fiber protein, gp68 (Supplementary Fig. S4).

A striking feature of the 80α baseplate is the presence of two tail fiber proteins, FibL and FibU, in addition to RBP. These proteins are probably involved in host interactions, although it is not known whether they bind to WTA or other components of the staphylococcal cell wall. Although FibL was essential for 80α infectivity, this was likely because deletion of ORF62 also caused loss of RBP (Supplementary Fig. S3). FibU was not essential, and deletion of ORF68 had no effect on either infectivity or baseplate integrity (Supplementary Fig. S3). The two C-terminal domains of FibL are similar to those of the tail fiber protein of phage K (Supplementary Fig. S4), a large-genome myovirus that infects *S. aureus* strains belonging to several lineages (*37*). Thus, while FibL and FibU may not be strictly required for infection of RN4220 under standard laboratory conditions, they could facilitate adsorption under certain conditions and be involved in binding divergent *S. aureus* strains that express altered WTA.

The presence of multiple copies of receptor binding complexes is a common theme among phages of Gram-positive bacteria. 80α has six trimers of RBP, while P68 has twelve (*36*). Lactococcal phage TP901-1 has 18 trimers of its receptor binding protein BppL (*22*). Presumably, the presence of multiple copies of the receptor binding proteins increases the avidity of the low-affinity interaction with WTA and surface polysaccharides. In contrast, phages like λ and SPP1 that use protein receptors—a high affinity interaction—generally do not have multiple receptor binding protein complexes.

Receptor binding is only the first step in the infection process that is mediated by the baseplate (Fig. 6A). Once bound, enzymatic activities associated with the baseplate are used to degrade the cell wall peptidoglycan and penetrate the cell wall. This process most likely requires conformational changes in the baseplate in order to expose the hydrolase domains of Tal and Hyd (Fig. 6B). We previously observed that treatment of 80α with heat or low pH led to clustering of baseplates, presumably due to exposure of hydrophobic sequences resulting from a conformational change (*16*). The observed conformational differences in the stem between the 80α and ϕ11 RBPs and the phage P68 tail fiber (Fig. 5B, C) suggest that the hinge might constitute a pivot point that allows the RBPs to rotate. This movement might assist in orienting the baseplate perpendicular to the cell wall. Straightening of the RBP stem might also lead to release of FibL, which is bound to the RBP hinge (Fig. 5D). Similarly, in p2, reorientation of its ORF18 receptor binding protein occurred as a consequence of a Ca^2+^-triggered conformational change in Dit (*25*).

**Figure 6.**
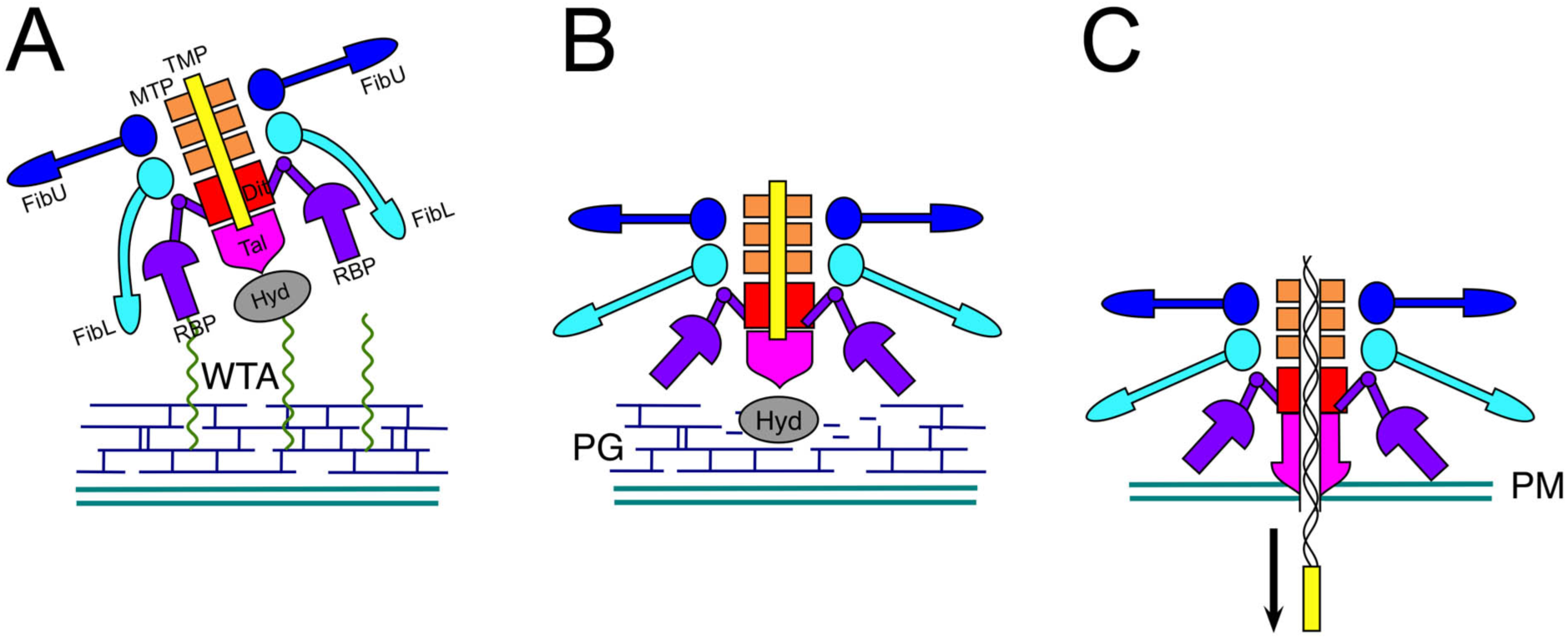
Model for conformational changes in the baseplate during the infection process. (A) Initial binding of RBP to WTA. FibL and FibU may bind as secondary receptor binding proteins to other surface structures. (B) Conformational changes in RBP lead to exposure of enzymatic activities associated with Hyd and Tal, allowing degradation of the cell wall peptidoglycan (PG). (C) Penetration of the plasma membrane (PM) by the Tal rod helices triggers release of TMP leading to ejection of DNA through the central channel.

Once the peptidoglycan layer is degraded and Tal reaches the plasma membrane, the α-helices in the rod domain may be involved in plasma membrane penetration, ultimately resulting in DNA ejection (Fig. 6C). This step has to be tightly regulated to avoid premature DNA ejection. How is plasma membrane penetration signaled to the capsid in order to initiate DNA release? Other studies have revealed no apparent conformational change in the tail tube associated with the infection process in siphoviruses (*31*). Even in the myoviruses, tail contraction itself is not the signal that leads to ejection (*11, 34*). Our observation of TMP α-helices associated with Tal resolves this conundrum. Conformational changes in Tal could release the TMP α-helices, which then might act as a “pull cord” that drags the DNA along with it as it is injected into the host (Fig. 6C).

Most temperate bacteriophages with long tails (myoviruses and siphoviruses) share common features in structural and genomic organization, reflecting their common evolutionary origin (*40*). According to the “modular theory” of phage evolution, phage genomes evolve by exchanging modular units consisting of protein domains, as opposed to whole genes or transcriptional units. In accordance with this principle, the tail and baseplate proteins in 80α are based on a set of conserved modules that is found in wide variety of phages. In 80α, MTP, Dit and Tal all incorporate the tail tube fold, one of the most highly conserved bacteriophages structural modules (*30*). In contrast, the peripheral structures are far less conserved between 80α and other phages, presumably reflecting different adsorption strategies between bacteriophages of different hosts. Nevertheless, the proteins that make up the peripheral structures share common domains with proteins from several bacteriophages, reflecting the modular evolution of these proteins. Modules such as Ig-like N-terminal domains, RBP tower domains and phage K tail fiber CTDs are repeated across phages from a broad range of hosts (Supplementary Fig. S4).

Our study provides a structural basis for understanding host specificity and the infection process in staphylococcal bacteriophages. Since bacteriophages are the primary mediators of horizontal gene transfer in staphylococci, this process is critical to the dissemination of virulence factors and the evolution of pathogenicity in *S. aureus* and other pathogens. The ability to infect a variety of strains and species will also be important for the development of effective therapeutic strategies utilizing phages.

## METHODS

### 1. Production of 80α tails

80α tails were produced by mitomycin C induction of *S. aureus* strain ST247, which is an 80α lysogen with a deletion of residues 1-13 from the scaffolding protein, gp46 (*16*). The tails were purified from the lysate by PEG precipitation and CsCl gradient purification, as previously described.

### 2. Electron microscopy

Cryo-EM samples were prepared on nickel Quantifoil R2/1 grids, as previously described (*15*). An initial cryo-EM data set consisting of 51 images was collected on SO-163 film on a Philips CM20 FEG microscope. The films were scanned with a Nikon 9000ED film scanner at 4000 dpi, corresponding to 1.27 Å/pixel, and used to generate the initial model for reconstruction. High-resolution data was collected on an FEI Titan Krios microscope equipped with a Direct Electron DE-20 detector at the SECM4 consortium at Florida State University. A smaller dataset of 715 micrographs and a larger dataset of 6,474 micrographs were collected at 29,000x magnification, corresponding to 1.21 Å/pixel in the specimen, and with typical defocus of 1.0–3.0 µm.

### 3. Three-dimensional reconstruction

An initial C6 starting model was generated from 4,180 particles picked from the film data, then used to reconstruct and refine 6,906 particles picked from the smaller FSU dataset, using EMAN 1.9 and EMAN 2.2 (*17*). The resulting 9.3 Å resolution (FSC_0.143_) reconstruction was in turn low-pass filtered and used as an initial model in the reconstruction of the larger FSU dataset in RELION-2.1 (*18*). 515,106 particles were picked from the 6,474 micrographs following frame alignment with dose weighting using MotionCor2 and CTF correction using Gctf, all from within RELION. After extensive 2D and 3D classification, the final reconstruction was calculated from 74,294 particles using the gold standard approach and assuming C6 symmetry throughout the reconstruction process. Following mask generation and map sharpening in RELION, the estimated resolution reached 3.75 Å (FSC_0.143_) (Fig. 1G and Supplementary Figs S1 and S2).

Generation of the C3 reconstruction used for modeling Tal and Dit required separation of the non-identical orientations of the C3 Tal trimer superimposed as a result of the initial assumption of C6 symmetry. The C6 reconstruction was segmented in UCSF Chimera (*41*) and a mask generated from all non-Tal density was used in subtracting the non-Tal density from the particle images using RELION-3.0 (*19*). The resulting signal-subtracted images were C6 symmetry-expanded using *relion_particle_symmetry_expand* to create three copies of each particle where the Tal trimer, regardless of initial orientation, was superimposed with the 60° rotated non-identical orientation. Masked 3D classification in RELION-3.0 of the expanded dataset using the Tal density from the C6 reconstruction as a reference yielded two good classes related by a 60° rotation and excluding particles with suboptimal Tal density. The more populated class was cleaned of duplicate particles, the orientations were transferred to the original, unsubtracted dataset, and the particles were reconstructed assuming C3 symmetry using RELION-3.0 with the C6 baseplate reconstruction as an initial model. Following post-processing, the resolution of the C3 reconstruction reached 4.07 Å resolution (FSC_0.143_, Fig. 1G). Focused reconstructions for RBP and the N-terminal parts of FibL and FibU were done in the same way, but without symmetry expansion and with the application of C6 symmetry to optimize the resolution of each for atomic model refinement. Reconstruction of signal-subtracted images lacking all non-FibL density (Fig. 5G) in RELION-3.0 allowed assembly of a full-length FibL pseudo-atomic model by rigid-body fitting matched structures from HHpred to the density. These models were manually mutated to the FibL sequence and joined together and to the refined FibL NTD structure (Fig. 5H). The FSC curves and ResMap analysis for the various reconstructions are shown in Supplementary Figs S1 and S2, respectively. The C6 and C3 maps were submitted to EMDB with ID numbers EMD-XXXX and EMD-YYYY, respectively.

### 4. Model building and refinement

The initial model for RBP was adapted from the ϕ11 RBP (PDB ID: 5EFV)(*23*). Initial models for the remaining proteins were generated using I-TASSER (*28*). Those best matching the baseplate reconstruction were fitted into the density using UCSF Chimera (*41*). Changes to the protein sequence, extension of the models, and manual adjustments to the protein backbone, including local real-space refinement with secondary structure, torsion angle, rotameric, and Ramachandran restraints, were made in *Coot* (*42*). The positions of large aromatic and basic side chains, as well as predicted secondary structures, were compared to the baseplate protein sequences and initial models to confirm the identity of proteins in the baseplate density and to correctly place the models during manual adjustment. The models were iteratively refined using real-space refinement in Phenix (*29*), followed by manual correction and local real-space refinement in *Coot* guided by geometry reports from MolProbity (*43*) and EMRinger (*44*). All models except Tal and TMP were initially refined into the C6 reconstruction. The final Dit, Tal and TMP models were refined in the C3 reconstructions, while RBP, FibU and FibL were further refined in their respective focused reconstructions. Simulated annealing, rigid body fitting, and morphing were included in early cycles of real-space refinement in Phenix. Secondary structure restraints and non-crystallographic symmetry restraints generated from the models by Phenix were included throughout. The model statistics for each protein are listed in Supplementary Table S1 and the quality of the fit (FSC) is shown in Supplementary Fig. S1. The refined models were submitted to the RSCB protein data bank with PDB ID ZZZZ.

## ACKNOWLEDGMENTS

We are grateful to John Spear and Duncan Sousa at Florida State University for assistance with the high-resolution data collection at the Southeastern Consortium for Microscopy of Macromolecular Machines (SECM4). We are also grateful to Dr. Alasdair C. Steven for access to the EM facility at The National Institute of Arthritis and Musculoskeletal and Skin Diseases (NIAMS), where the initial data collection was carried out, to Cynthia M. Rodenburg for technical assistance at the UAB cryo-EM facility, to Dr. José Penadés for providing the 80α baseplate deletion mutants, and to Dr. Pavel Plevka for sharing the coordinates for phage P68. Access to SECM4 was provided through National Institutes of Health (NIH) grant U24 GM116788 to Dr. Kenneth A. Taylor at FSU. ADD was supported by the intramural research programs of the National Institute of Allergy and infectious Diseases and NIAMS.

This project was supported by NIH grant R01 AI083255 to T.D.

## AUTHOR CONTRIBUTIONS

J.L.K. carried out the majority of the data processing, modeling and analysis; K.A.M. produced the phage material; A.D.D. carried out the initial cryo-EM; T.D. directed the research and data analysis and performed some of the data processing. All authors were involved in writing and editing the manuscript.

## SUPPORTING INFORMATION

**Supplementary Figure S1.**
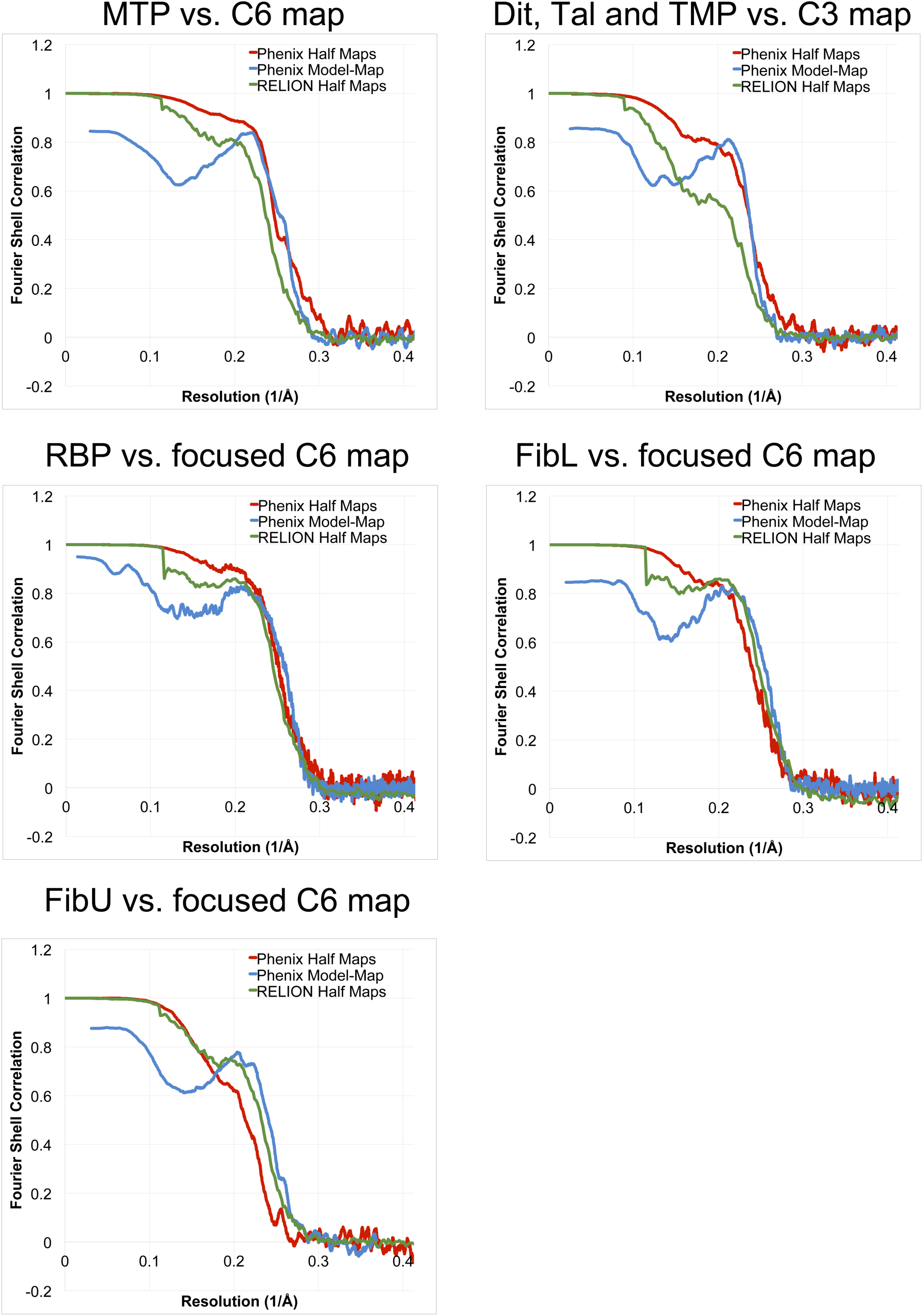
Fourier Shell Correlation (FSC) curves for the various model/map combinations. For each refinement, the gold-standard half-map FSC plots for the appropriate maps, calculated in RELION (green curve) and in Phenix (red curve) are shown. The blue curve represents the corresponding model-to-map FSC plot, calculated in Phenix.

**Supplementary Figure S2.**
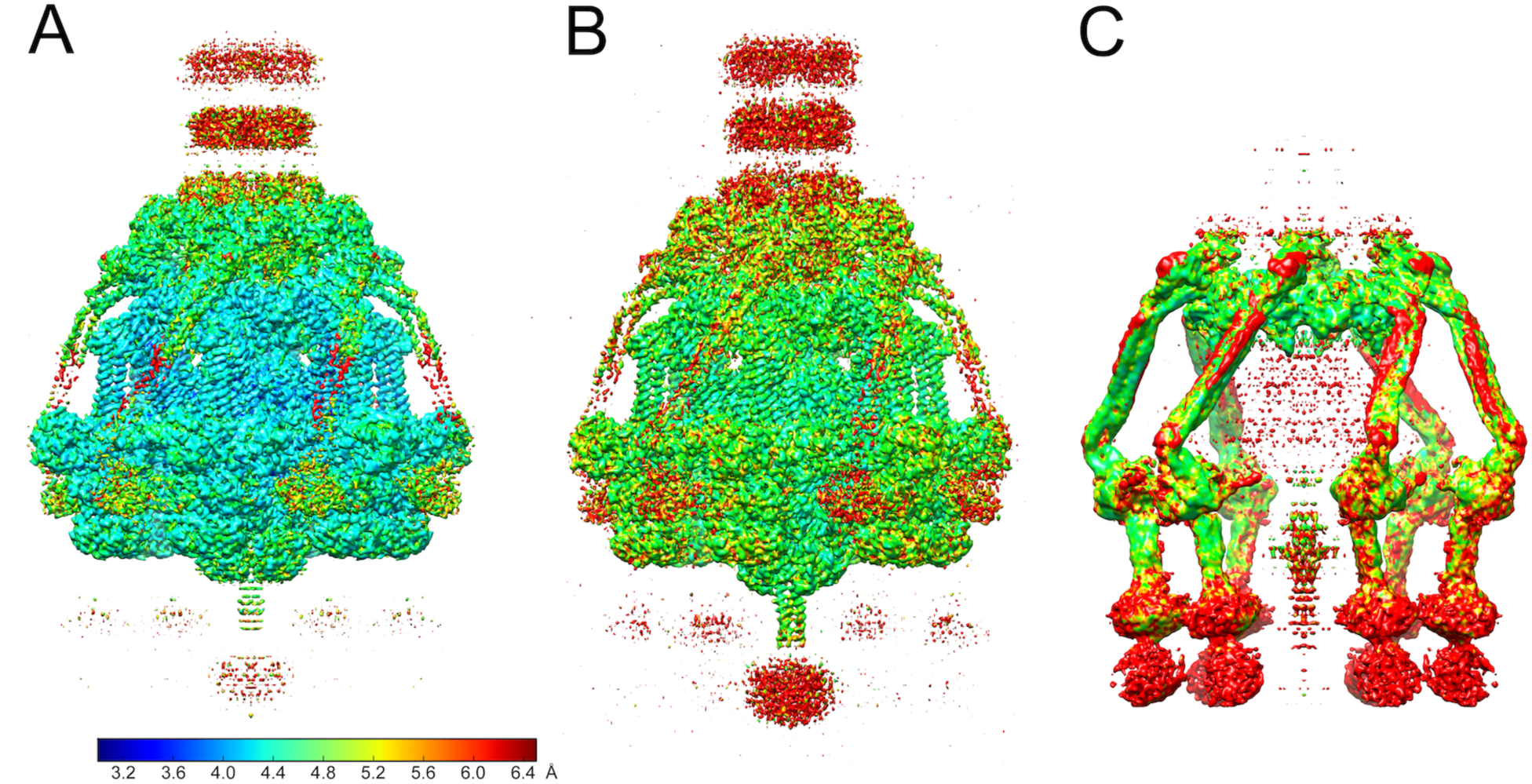
ResMap analysis for the C6 symmetrized map (A), the C3 map (B) and the C6 signal-subtracted map used to model FibL. Map resolution is shown from 3.0 Å (blue) to 6.5 Å (red) according to the color bar.

**Supplementary Figure S3.**
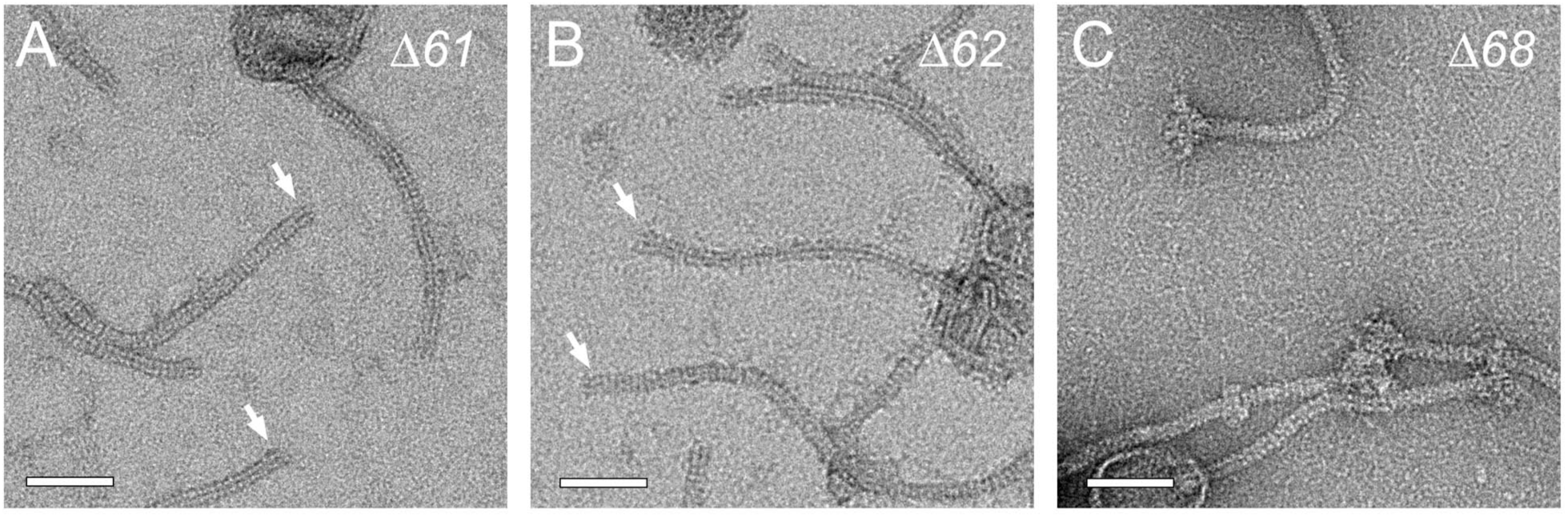
Baseplate deletions. Phage particles produced by 80α lysogens with deletions of (A) *ORF61* (RBP), (B) *ORF62* (FibL), and (C) *ORF68* (FibU). The arrowheads point to baseplates missing peripheral structures in (A) and (B). Scale bars = 50 nm.

**Supplementary Figure S4.**
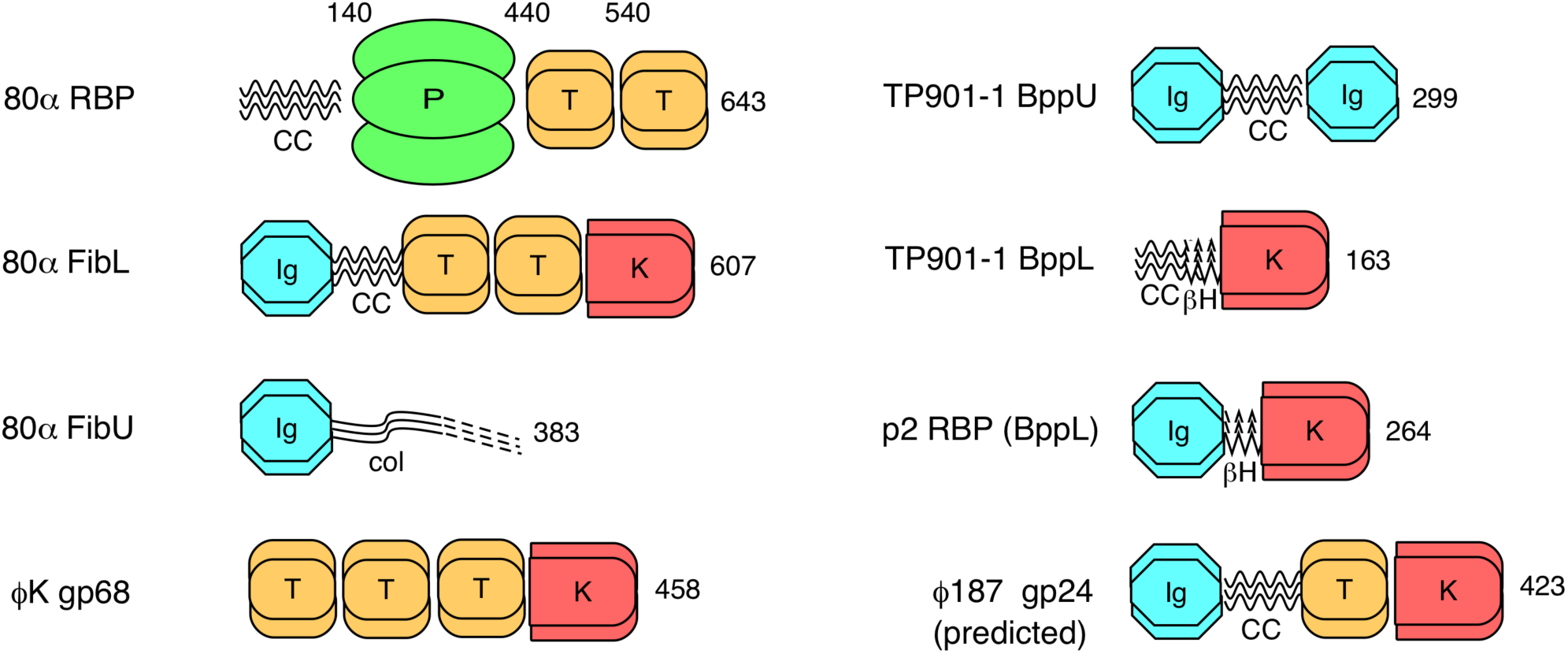
Modular domain organization of 80α baseplate proteins RBP, FibL and FibU compared to tail fiber/receptor binding proteins from TP901-1, p2 and phage K (crystal structures). The predicted structure of the ϕ187 tail fiber is also shown. Domains are labeled: CC, coiled coil; P, β-propeller platform domain; T, tower domain; K, phage K gp68 CTD; Ig, immunoglobulin-like domain; βH, beta-helix; col, collagen-like helix. Protein length and relevant residue numbers are indicated.

**Supplementary Table S1.**
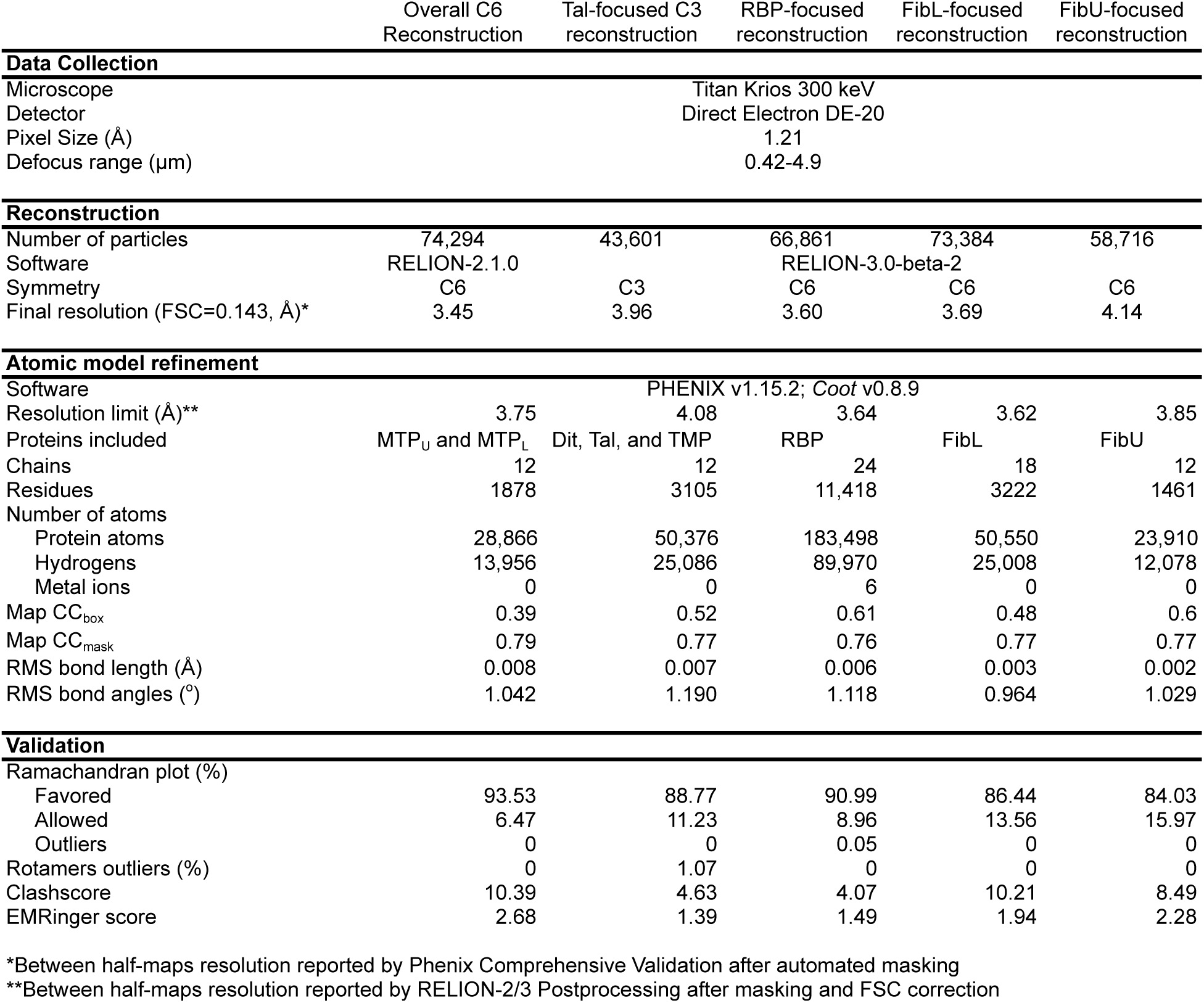
Reconstruction and modeling statistics.

